# Automatic Generation of Model Sequences for Complex Regions in Assembly Graphs

**DOI:** 10.64898/2026.03.06.710180

**Authors:** Dmitry Antipov, Ying Chen, Marco Sollitto, Adam M. Phillippy, Giulio Formenti, Sergey Koren

## Abstract

Recent developments in genome sequencing and assembly technologies have enabled the automated assembly of vertebrate chromosomes from telomere to telomere. However, for some long, highly similar repeats, genome assemblers may lack sufficient information to unambiguously resolve the sequence, leaving tangles in the assembly graph and gaps in the final assembly. In recently published genomes, such gaps are often closed by manual graph curation, a process that is labor-intensive, error-prone, and sometimes infeasible. This can leave important genomic repeats, such as recently duplicated genes, misassembled or excluded from the final assembly. Here we present the Trivial Tangle Traverser (TTT) algorithm that finds optimized resolutions of assembly graph tangles. TTT uses depth of coverage and read-to-graph alignment information in a two-stage process to identify evidence-based traversals that are consistent with the underlying data. First, sequence multiplicities are estimated through mixed-integer linear programming, after which an Eulerian path is found in the derived multigraph and optimized through a gradient-descent-like approach. We evaluate TTT traversals on the HG002 human reference genome and demonstrate its use to characterize a previously unassembled amplified gene array in the zebra finch genome.

**Availability:** TTT is available at https://github.com/marbl/TTT

## Introduction

Advances in sequencing technologies have enabled a new goal for de novo genome assembly: the complete assembly of entire chromosomes from “telomere to telomere” (T2T). In the T2T-CHM13 project (Nurk et al. 2022), assembly continuity was achieved by combining previous-era genome assemblers along with specialized semi-manual processing. Subsequently, specific tools focusing on automated T2T genome assembly were developed (Cheng et al. 2021; Garg et al. 2021; Rautiainen et al. 2023; Cheng et al. 2024; Lorig-Roach et al. 2024; Antipov et al. 2025). However, despite all the progress in sequencing technologies and assembly algorithms, some complex genomic regions are still unresolvable with the current read length and error profiles (Li and Durbin 2024; Rautiainen 2024).

T2T assembly projects benefit from either parental (Koren et al. 2018; Rautiainen et al. 2023) or long-range (Cheng et al. 2021; Garg et al. 2021; Antipov et al. 2025) sequencing data for haplotype separation and scaffolding. Thus, T2T genome assemblers can “jump over” complex genomic regions, leaving a gap between contigs in a scaffold, rather than just stopping at those regions. This can lead to significant differences between the number of T2T contigs and scaffolds in the assembler’s output. For example, the verkko2 paper reported a median count of 21 T2T contigs versus 39 T2T scaffolds across 17 HPRC human genome assemblies (Antipov et al. 2025).

T2T assembly projects (Nurk et al. 2022; Yoo et al. 2025; Olagunju et al. 2025; Garg et al. 2024; Miga and Eichler 2023) aim to resolve gaps and close them by postprocessing the results of the assembler software. Most commonly used gap-closing tools like LR_Gapcloser, TGS-GapCloser, and SAMBA (Xu et al. 2019, 2020; Zimin and Salzberg 2022) rely on finding individual reads that bridge the gap. This limits their usefulness to closing relatively short gaps (100–200 kbp). DEGAP (Huang et al. 2024) can fill larger gaps through an iterative extension strategy, stopping when no extension is available or there is sufficient similarity between the extension and the gap boundary. However, these tools can leave gaps in complex regions, such as large repetitive arrays and segmental duplications (Zhao et al. 2024). Worse, this can lead to gaps being filled with incorrect sequences that can escape validation check due to read mapping ambiguity in repetitive regions (Chu et al. 2025) and may go unnoticed by future users of the reference assembly. T2T projects therefore often rely on semi-manual inspection of the assembly graph followed by manual path selection using project-specific scripts (Nurk et al. 2022; Kim et al. 2025). The benefit of this approach is that it generally produces a resolution that is consistent with the underlying sequencing data (represented by the assembly graph), but this process requires domain expertise, is labor intensive, and lacks the reproducibility of automated tools.

Here we present TTT, a tool that replaces the error-prone manual gap closing process with the identification of an optimal tangle traversal in the assembly graph that leads to the automated construction of a patch sequence. The problem of approximate gap closure is different from a typical local or global de novo genome assembly problem. Most genome assembly tools prioritize correctness, and so usually stop in ambiguous regions, providing a gapped or fragmented assembly. In complex regions, these highly fragmented or incomplete sequence assemblies can lead to missing gene duplications, hindering analysis. To reveal these dark regions, TTT prioritizes completeness and aims to find a traversal which is most consistent with the input data. Instead of providing a partial solution (i.e. a set of fragmented contigs), it always produces a single continuous path through the tangle that is most consistent with the sequencing data. Therefore, in the case of multiple equally good solutions (e.g. in the case of an exact repeat larger than read length) it will select one of the paths even if its correctness cannot be unambiguously verified. Given current sequencing read lengths and accuracies, there can be multiple such regions in large, complex genomes that are essentially unsolvable. Thus, we refer to the output of our tangle traversals as “model sequences” rather than “assemblies” to communicate this uncertainty.

Below we describe the algorithm in detail, compare TTT with a general-purpose assembler on a large human genome reference test set, and highlight its performance on two previously unassembled megabase-scale gaps on the Z chromosome of the zebra finch genome, enabling the study of this complex ampliconic gene array for the first time.

## Results

To close assembly gaps of arbitrary size we have implemented a method named the Trivial Tangle Traverser (TTT). TTT takes as an input an assembly graph, coverage information and read alignments and searches for a path through the corresponding region of the assembly graph which has the highest support from sequencing coverage and read alignments. **Figure 1** shows a simplified example of TTT’s input and output. TTT relies on an assumption of sequencing coverage uniformity to estimate the multiplicities of graph edges within a tangle. Since each gap is considered separately, uniformity conditions are only required for local regions and not across the whole genome, which mitigates the effects of localized sequencing coverage biases, such as within tandem satellite arrays. TTT reduces the multiplicity estimation problem to a mixed-integer linear programming problem (Meyer 1974, 1975), a well-studied class of optimization problems that can be addressed using a variety of highly optimized heuristic solvers. Then, with the determined edge multiplicities and corresponding multigraph, TTT searches for an Eulerian path with the best support from read-to-graph alignments through a gradient-descent-like optimization. Details of the TTT algorithm are described in the Methods and Supplementary Materials.

**Figure 1.**
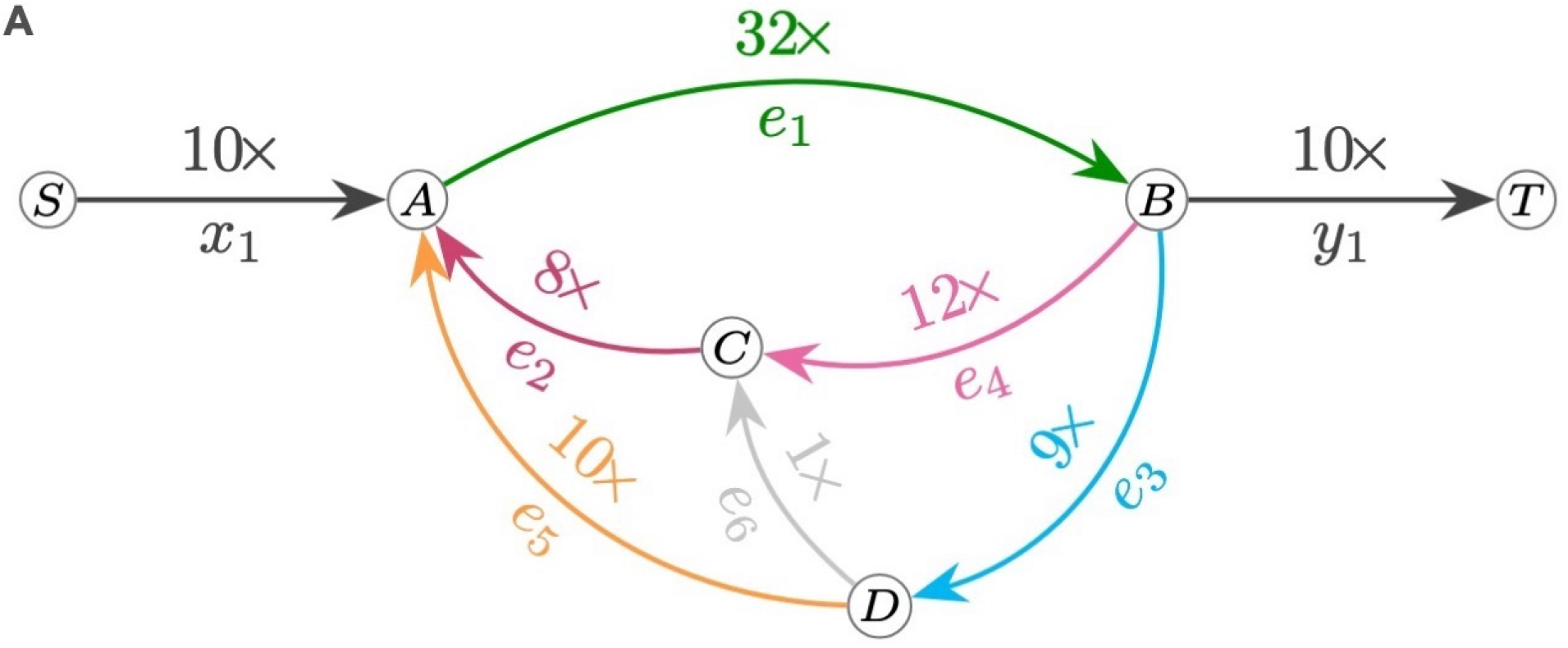
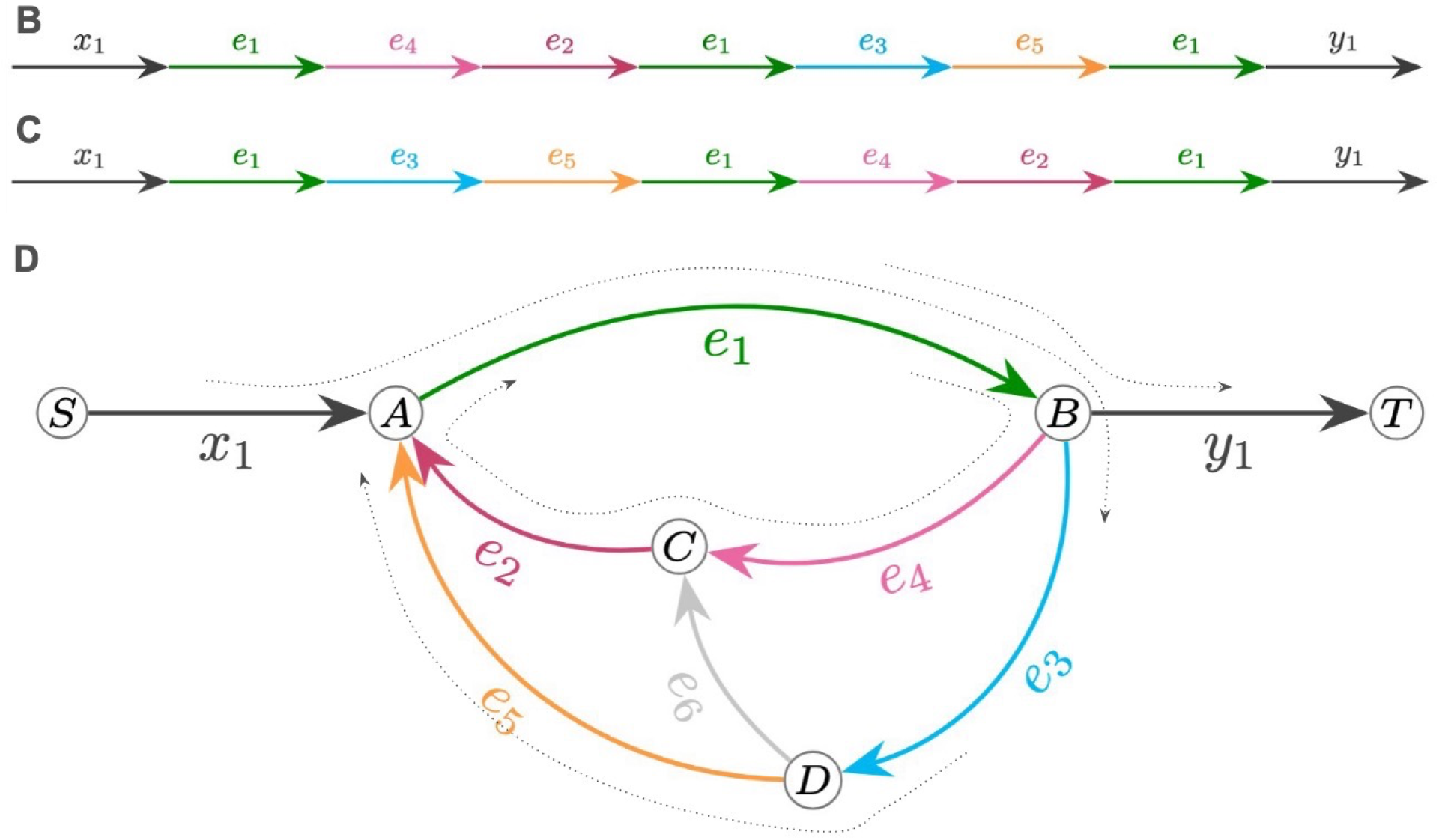
A) Example of an assembly graph tangle with boundary edges x_1_ and y_1_ and tangle edges e_1_ e_2_…e_6_ additionally labeled with coverage information. Capital letters A,B,C,D,S,T represent the graph vertices. B) and C) Two possible underlying genomic traversals consistent with the coverage data. D) Read alignments to the same graph (dotted curves) demonstrates that the traversal in panel C has better support.

### Reference-based evaluation using T2T-HG002

The overarching goal of TTT is to provide plausible sequence models for residual tangles that conventional graph traversal algorithms fail to resolve. To test the ability of TTT to generate relevant traversals, we first extracted all tangles from a verkko assembly of the HG002 sample and compared TTT with verkko’s MBG-based repeat resolution (Rautiainen et al. 2023) on the same tangles, as well as with the HG002 (Hansen et al. 2025) reference sequence.

For evaluation, we used a HiFi-only graph, which allows us to compare against verkko’s ONT-based repeat resolution on exactly the same tangles. We defined tangles in the graph based solely on the graph structure (see **Supplementary Methods 2** for details). We filtered out all tangles that had neither two nor four boundary nodes, as well as tangles with four boundary nodes that were not resolved or scaffolded by verkko (for which we cannot infer boundary-node matching from verkko’s scaffolds). On the remaining 220 tangles, we ran TTT with all parameters set to their default values. Two runs crashed, 218 succeeded, providing 397 traversal paths (179 tangles were traversed by two genomic paths).

We compared TTT’s traversals with those inferred from the verkko 2.2.1 results for the same tangles. In most cases (363 of 397) they were identical. In three cases detected tangle borders didn’t belong to the same verkko paths. There were five cases where we cannot compare TTT against verkko due to gaps in the final verkko’s contigs, and one case where verkko closed the gap in the assembly graph by adding sequence from ONT reads (an option not available to TTT). For the remaining 25 cases where verkko and TTT traversals differed, we compared them by aligning corresponding sequences to HG002 v1.1 reference with minimap2 (Li 2018) using minimap’s alignment score of the best alignment as the measure of quality (see Table S1). In eight cases, TTT’s resolution scored better than verkko’s, while in 17 it was worse. We manually verified all cases where TTT’s score differed substantially with verkko’s and confirmed that all those cases were because of uneven ONT coverage, which led to errors in multiplicity estimation. For each tangle, the final alignment-based score is the same for all (10) different random initial Eulerian paths. Thus, we suppose that it is highly unlikely (although possible) that these scores and final paths correspond to local but not global optima.

This test focused only on those tangles where a verkko resolution was available and therefore is biased towards simpler, resolvable tangles. While this does not demonstrate an improvement over verkko’s resolution, the test confirms that TTT is consistent and comparable with existing resolution methods and serves as a sanity check.

For additional evaluation we defined two reference-free metrics for internal evaluation of TTT’s results. The first is the divergence between multiplicities and coverage, normalized by coverage and number of edges, 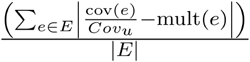. The second is the fraction of read alignments corresponding to the tangle that are not supported by the TTT traversal (i.e. are not exact substrings in the alphabet of edge labels). **Supplemental Figure S5** shows those metrics on all tangles. Problematic tangles tend to have higher values for both of these metrics.

### Novel tangle resolution in the zebra finch near T2T genome

Given the success of our reference-based comparison, we set to test TTT by resolving the remaining gaps in a recently published near T2T zebra finch (*Taeniopygia guttata*) genome (Formenti et al. 2025). The current reference (GCA_048771995.1) is based on verkko 2.2.1 (Antipov et al. 2025) assembly. The assembler’s output contained 4 gaps on the chrZ scaffold associated with highly collapsed repetitive sequences flanking the gaps (**Figure 2A, Supplementary Figure S3A**). Two of these tangles were resolved using standard approaches, that is by manual graph curation and by patching using a different Verkko assembly that was able to span the gap.

**Figure 2.**
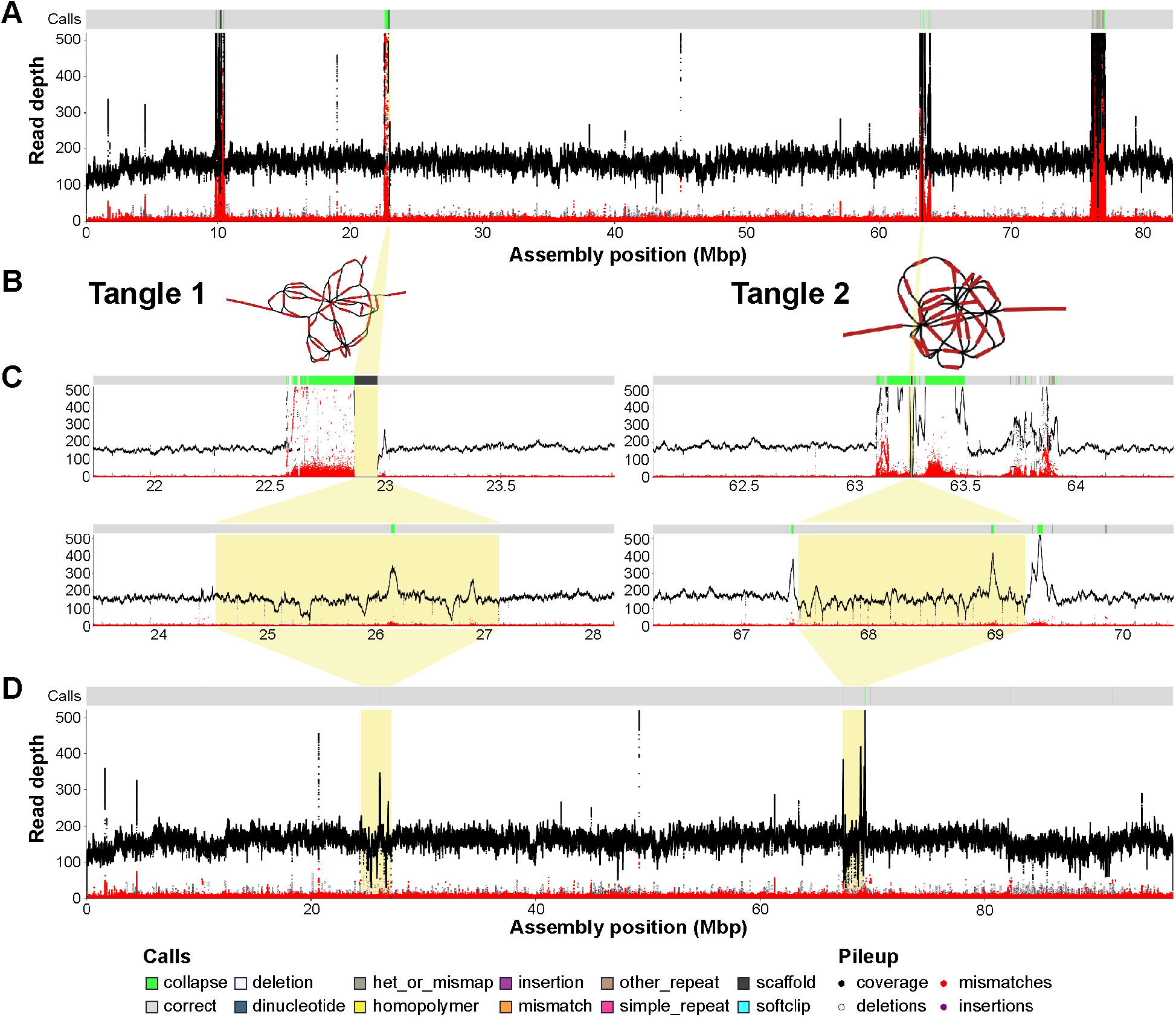
NucFlag ONT coverage plots for (A) zebra finch chromosome Z before resolving the gaps (NCBI accession NC_133063.1), two tangle regions (B) before and after running TTT (C), and (D) zebra finch chromosome Z after resolving the four gaps. The start and end position for the plotting region in (B) and (C) are the same for each tangle, with the tangles highlighted in yellow. The primary read coverage is shown in black while secondary allele frequency is shown in red. Peaks or drops in coverage (black line) indicate assembly issues, either collapse (peaks) or mis-joins (drops). However, some variation is expected due to coverage biases in sequencing data. Peaks in the secondary allele (red) support classifying a collapse as this indicates a large subset of reads disagree with the assembly, likely because these reads belong to a different repeat copy. **Supplementary Figure S3** shows results using Oxford Nanopore sequencing. The bar above each coverage plot indicates automatically annotated errors in the assembly, colored by error type. The highlighted regions show a dramatic decrease after TTT resolution (C).

Two residual tangles were considered to be too complex for manual curation by the zebra finch T2T project and motivated TTT development (**Figure 2B**). TTT was run on these two unresolved tangles in the Verkko’s assembly. The first tangle contained 45 edges, and the second tangle contained 32 edges In both cases, the tangles were modeled with TTT and then the sequence was incorporated in the existing reference (2.6 Mbp and 1.8 Mbp for the two tangles, respectively). Validation using NucFlag of the original assembly and the final gap-filled result showed a dramatic reduction in read pileups and secondary allele frequency (**Figure 2C, Supplementary Figure S3B & S3C**). DEGAP filled the same two gaps but the sequence was shorter than TTT (0.4 Mbp and 0.1 Mbp) and NucFlag flagged the resolved regions as collapsed (**Supplementary Figure S4**).

The TTT reconstruction enables the study of this region of zebra finch chrZ for the first time (**Figure 3**). Gene annotation of the whole chromosome with miniprot shows multiple clusters of ampliconic gene arrays dominated by p21-activated serine/threonine kinase 3-like genes (*PAK3L*) sequences (**Figure 3A**). Strikingly, all 4 original gaps coincide with *PAK3L* ampliconic gene arrays. Focussing on the regions resolved by TTT, the first tangle contains 12 units of median length 268 kbp and the second contains 35 units of median length 59 kbp, well beyond the resolution potential of bridging-read-based gap-closing tools (**Figure 3B,C**). Miniprot annotations of *PAK3L* as queries reveal exactly 200 copies of the *PAK3L* gene organized in 10 main clusters (**Figure 3A**). Of the 200 copies, only 4 of them present frameshift issues, suggesting that they may largely be functional

**Figure 3.**
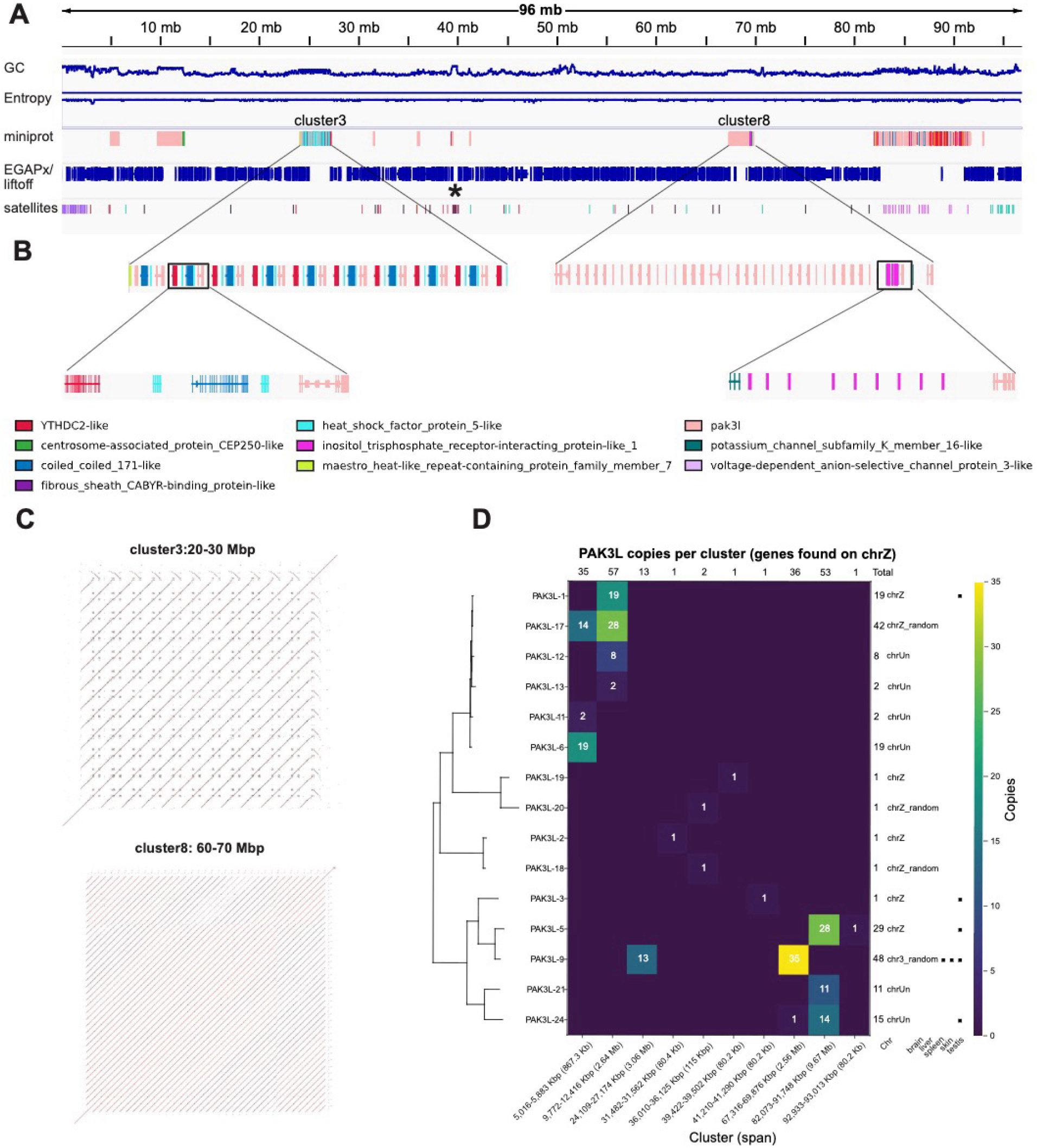
A) Integrative Genomics Viewer (IGV) representation of chromosome Z after gap resolution. Tracks include GC content, sequence entropy, miniprot alignments using as query the proteins identified by EGAPx gene annotations, Liftoff-transferred EGAPx gene annotations from previous T2T assembly for chrZ, and tandem repeat annotations. The centromere is indicated by an asterisk. Ten different clusters of the *PAK3L* genes are visible in the miniprot track B) Overview of the cluster 3 and 8 regions (gaps filled by TTT), with an inset highlighting the duplicated gene structure. Cluster 8 shows the duplication of the *PAK3L* gene and other genes, such as the inositol trisphosphate receptor-interacting protein-like 1 gene. C) Dot plots of the two clusters. D) Heatmap showing the copy number of each PAK3L gene across clusters, together with a schematic summary of the genes for previously reported chromosomal localization and tissue-specific expression (Kong et al. 2010).

The *PAK3L* ampliconic gene arrays are of particular interest for the zebra finch biology, as 31 *PAK3L* genes were previously annotated in the zebra finch genome, with 11 copies associated with chrZ (Warren et al. 2010), of which 5 were assigned to coordinates while 6 were found on unplaced sequences (Kong et al. 2010). Another 17 copies could not be assigned to any chromosome. Reported sequence similarity was high, and the duplications were all considered to have occurred within the past 5 Mya. When the gene predictions are assigned to the best *PAK3L* gene given their sequence homology, they reveal a pattern in which specific *PAK3L* genes are associated with different clusters (**Figure 3D**). For instance, the first cluster is using only *PAK3L-6, PAK3L-11*, and *PAK3L-17*. A clear tendency for more similar gene copies to colocalize is also visible throughout the chromosome. Importantly, the two initially unresolved tangles that were modelled by TTT share the same *PAK3L-9* gene in multiple copies, with the second tangle having the highest number of the same *PAK3L* genes in the whole chromosome. This may explain why these regions were particularly recalcitrant to assembly, while the regions with the two clusters with the largest numbers of copies overall were more diverse and did not require sequence modeling with TTT.

*PAK3L* genes in different clusters also appear to have evolved a different intronic content and other cluster-specific gene duplications are sometimes present (**Figure 3B**), which may explain why they appeared as completely distinct in the assembly graph. These other duplications include YTHDC2-like, heat shock factor protein 5-like, and coiled-coiled 171-like genes in cluster 3, as well as inositol triphosphate receptor interacting protein-like genes and potassium channel subfamily K member 16-like genes on cluster 8. Interestingly, in contrast to previous reports (Wirthlin et al. 2014) that described songbird *YTHDC2* as highly truncated and *YTHDC2L1* as retaining only the HA2 and OB-fold domains, the genes identified here contained all the functional domains (DEAD, HA2, Helicase_C, OB-fold, R3H, and YTH). Relative to *PAK3*, which resides on chr 4A, *PAK3L* paralogs were shown to lack the 5’ exons encoding the N-terminal regulatory domain (Kong et al. 2010). Yet for at least 7 of them, evidence of expression, particularly in the testis but also in the brain (*PAK3* expression is considered restricted to the central nervous system) (**Figure 3D**), and of positive selection on 9 of them, was previously reported (Kong et al. 2010). Interestingly, *PAK3L-9*, i.e. the copy present in the two tangles resolved by TTT, was reported to be expressed in multiple tissues (testis, spleen, and skin). While the function of *PAK3L* remains to be elucidated, it is likely to contribute to songbird-specific biology. The resolution of the zebra finch chrZ clearly shows how TTT enabled the study of this biology at chromosome resolution.

## Discussion

Here we described a method for constructing model sequences for the most complex genomic regions that are left unresolved even with the state of the art T2T assemblers and sequencing technologies. TTT searches for the complete traversal that best fits the input data, and not the partial but more conservative outputs of most genome assemblers. Separating the edge multiplicity determination problem and the search for the best traversal itself allows us to find an approximate solution for both using common optimization techniques—mixed integer linear programming and gradient descent optimization.

We have shown that our evaluation metrics allow us to assign an a priori score or confidence measure that is consistent with reference-based evaluation. However, it remains unclear whether these metrics should be interpreted as an evaluation of the tangle-traversing sequence itself or merely of the intrinsic difficulty of the problem. The coverage-based metric clearly depends on coverage uniformity, whereas the read-based metric depends on tangle complexity and the number of graph edges involved. Consequently, we cannot provide any recommendation for metric values that could be used as thresholds separating plausible from non-plausible traversals.

The described algorithms can be extended to incorporate multiple sequencing technologies. One clear way to improve TTT’s traversal is to use coverage information from different platforms simultaneously when determining multiplicities, or to modify the scoring function to account for Hi-C or trio-based information. Another clear place for improvement is the support for constructing model sequences for multi-chromosomal repeats, which can be done in the similar way to the two-haplotype repeats resolution (described in **Supplementary Methods S1**).

TTT relies on coverage information for copy number estimation. However, some repeat arrays are known to exhibit significant sequencing coverage bias (Nurk et al. 2022), and we thus expect that the copy number may occasionally be mis-estimated. Furthermore, coverage gaps or significant errors in the assembly graph may preclude a traversing path, leaving those gaps unresolved. Nonetheless, TTT’s algorithm is designed to ensure that every edge of reasonable coverage is included in the final traversal, meaning all repeat variations will be present, although potentially arranged in the wrong order. We believe this provides a better representation of a repetitive region than a gap (missing data) or a collection of short contigs (difficult to analyze with typical sequence analysis tools).

Although plausible and difficult to refute by design, we suggest that TTT traversals should be considered model sequences of a priori lower quality than standard assembler output. This distinction can be represented in the final assembly in different ways—through quality scores, annotation, or simply by separating the model sequence from the assembled scaffold in a FASTA file. We emphasize that such a distinction should be represented *somehow*, and therefore we intentionally do not provide automatically “patched” T2T scaffolds as output so that users must make a conscious decision on how to incorporate the traversals.

We have described an algorithmic approach that is expected to close some of the remaining gaps in T2T scaffolds. Currently, TTT cannot be used to model inter-chromosomal repeats (e.g. rDNA tangles in human genomes), and it will not help in cases where gaps are caused by reduced coverage rather than complex repetitive structure. Although not applicable to all gaps in a genome assembly, we believe it represents an important step toward truly telomere-to-telomere genomes and the study of complex regions of the genome that are otherwise not represented in the assemblies and therefore not accessible to investigation.

By closing the remaining gaps of zebra finch chrZ using TTT, we were able to precisely map multiple ampliconic gene arrays, assigning distinct copies of the duplicated genes to their genomic loci on chrZ while also quantifying them. Zebra finch chrZ harbors large chromosomal inversions (Itoh et al. 2011) that have been shown to be strikingly associated with phenotypic variation in siring success (Knief et al. 2016; Kim et al. 2017; Knief et al. 2017). It did not escape our attention that such inversions appear to colocalize with the ampliconic gene arrays, and that the genes involved tend to be expressed in testis (Kong et al. 2010; Viitaniemi et al. 2024). Further research will be needed to precisely characterize these inversions at the sequence level and determine their potential phenotypic impact.

## Software availability

TTT is available on github (https://github.com/marbl/TTT). Version tagged v0.1.1 was used for benchmarking in this study.

## Data sets availability

No new data was generated for this study. Sequencing data for the HG002 dataset is available at HPRC’s Amazon cloud at: https://s3-us-west-2.amazonaws.com/human-pangenomics/index.html

Raw sequencing data for zebra finch is available in the NCBI SRA under BioProject accession PRJNA1241433. The fully resolved sequence for chrZ is available at: https://genomeark.s3.amazonaws.com/index.html?prefix=species/Taeniopygia_guttata/bTaeGut7/assembly_verkko_0.1/patching/

An accessioned version will be made available in NCBI upon publication.

## Funding

This research was supported by the Intramural Research Program of the National Institutes of Health (NIH) and by the National Human Genome Research Institute of the National Institutes of Health under Award Number R03HG013362. The contributions of the NIH author(s) are considered Works of the United States Government. The findings and conclusions presented in this paper are those of the author(s) and do not necessarily reflect the views of the NIH or the U.S. Department of Health and Human Services. This work utilized the computational resources of the NIH HPC Biowulf cluster (https://hpc.nih.gov).

## Acknowledgements

Authors thank Arang Rhie, Nancy Hansen, Juhyun Kim and Pavel Pevzner for helpful discussions, Ksenia Krasheninnikova, Ben Rosen, Sarah Fumagalli, Brendan Pinto, Jiadong Lin, and Jeffrey Schoenebeck for testing early versions of TTT and providing valuable feedback on its performance, and Keisuke K. Oshima for his help on NucFlag analyses.

## Author contribution

D.A. developed the TTT algorithm and wrote the initial manuscript draft. G.F., Y.C., and M.S. performed analysis of zebra finch repeat arrays. D.A., Y.C., M.S., G.F. generated figures. A.M.P. and G.F. provided project supervision and coordination. G.F. formulated the tangle traversing problem and supervised analysis of zebra finch repeat arrays. S.K. supervised development of the TTT algorithm and implementation. All authors edited the manuscript. All authors read and approved the final manuscript.

## Methods

### TTT algorithm

TTT considers tangle traversal as two independent sub-problems: (i) determining the multiplicities of tangle edges in the traversing path, and (ii) finding a path that is consistent with the largest number of read-to-graph alignments while respecting the edge multiplicities obtained in the first step. In the following we use the assembly graph representation with sequence stored on *edges* of the directed assembly graph (Chikhi et al. 2015). Every scaffold can be represented by a sequence of paths in the assembly graph separated by gaps. Although the most commonly used assembly graph representation (gfa, (Li 2016)) assumes that sequence is stored in nodes, a multiplex DBG (Bankevich et al. 2022) in gfa format can be easily transformed into a graph with sequence on edges by considering junctions as vertices and nodes as edges.

To formally define a tangle we use an approach similar to the definition of a snarl in (Chang et al. 2020). We say that *n* edge pairs (x_i_, y_i_) define a tangle in the assembly graph if: (a) all those edges belong to the same connected component C in the assembly graph (considered as undirected graph) (b) removal of those edges splits component C into multiple connected components C_1_, …C_k_ (c) one of those connected components C_t_ is significantly smaller than every other C_i_ (d) the end of each x_i_ and the start of each y_i_ belongs to C_t_. We call C_t_ a *tangle*; all edges in C_t_ are referred to as *tangle edges*, and the pairs (x_i_, y_i_) are referred to as *(incoming, outgoing) boundary* edges. **Figure 1** shows an example of a simple tangle and underlying genomic traversal.

TTT has an additional requirement for the tangles it resolves: every boundary edge x_i_ and y_i_ should be *unique* – i.e., non-repetitive and non-homozygous edges that should be present only once in a correct genome assembly. Since the correct assembly is unknown, in practice unique edges are identified based on their length and coverage. Informally, x_i_ can be considered as the last unique edge of the scaffold path before the gap, and y_i_ as the first unique edge after the gap. In contrast to snarls we must consider multiple pairs of boundary edges because a tangle edge can belong to multiple genomic paths; a common example is a repeat array shared by both haplotypes in a diploid genome.

First, we describe a simplified version of the algorithm for the tangles with one incoming and one outgoing boundary edge. For simplicity, we also ignore reverse complementation. Thus, we assume that for each edge *e*, only one of *e* and *reverse_complement(e)* is reachable from the tangle boundary edges, and we do not need to distinguish multiplicities of *e* and *reverse_complement(e)*. Changes required to process two-haplotype tangles and the case of inverted repeats are described in the **Supplemental Methods**.

### Identifying multiplicities of edges in the traversing path

In this step, we define optimal assignment of edge multiplicities according to the assembly graph structure and edge coverages. The primary requirement for the multiplicities is consistency with the coverage information; however, we also aim to ensure the existence of a path traversing the tangle with the assigned multiplicities, and that the resulting traversal does not discard edges with reasonable coverage.

TTT assumes that the coverage of traversals within the tangle is somehow uniform; otherwise, reliable estimation of edge multiplicities based on edge coverages would not be possible. All sequencing technologies are known to have some coverage bias (Rieber et al. 2013; Hon et al. 2020; Delahaye and Nicolas 2021; Carvalho et al. 2026), and thus global coverage uniformity across the genome is unlikely. However, we need only the local coverage uniformity, which is way more likely to observe. Moreover, with coverage information from multiple sequencing technologies available(which is a common situation for T2T assembly projects), TTT can select the one that is less biased.

We assume that each genomic path through the tangle has the same coverage, denoted later as Cov_u_. The initial value for this coverage is either provided by the user or estimated as the median coverage of unique boundary edges incident to the tangle, and it can be further tuned by the optimization process.

Comparing coverage information for repetitive edges with the coverage of known unique (haploid and non-repetitive) genomic regions can provide a straightforward estimate of edge multiplicity by dividing the edge coverage by Cov_u_ and rounding to the nearest integer. However, due to the coverage variation, such approximations may be impossible to satisfy – for example, at a vertex with two incoming edges and one outgoing edge, all assigned multiplicity one. Therefore, in addition to encouraging coverage-based multiplicities to be as close as possible to Cov_edge_/Cov_u_, we impose a natural set of flow conservation constraints: for each vertex, the sum of multiplicities of incoming edges is expected to equal the sum of multiplicities of outgoing edges.

Lastly, to ensure completeness of the traversal, we require that all reasonably covered edges (defined as edges with coverage at least 0.5 * Cov_u_) be included in the optimal traversal path at least once. Although this definition of “reasonably covered” is somewhat arbitrary, it is based on the assumption that an edge with substantial coverage is not “invented” by the assembler and therefore should not be excluded from the optimal traversal. Summarizing, with unknown integer edge multiplicities, we have a set of linear equations:

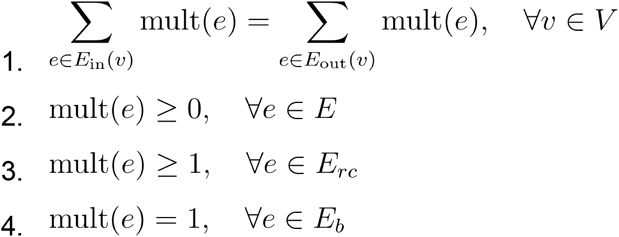

With an objective to minimize the difference between the edge multiplicities and their coverage-based estimates:

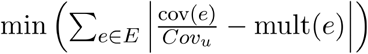

Where:

Mult(*e*) is the integer multiplicity of edge *e*.

cov(*e*) is the coverage of edge *e*.

*Cov*_*u*_ is the unique traversal coverage.

*V* is the set of all vertices in the tangle.

*E* is the set of all edges in the tangle.

*E*_in_ (*v*) and *E*_out_ (*v*) are the sets of incoming and outgoing edges for vertex *v*.

*E*_rc_ is the set of reasonably covered edges.

*E*_b_ is the set of boundary edges.

This is a mixed-integer linear programming (Meyer 1974, 1975) problem formulation to determine integer multiplicities for the edges. Although MILP is known to be NP-complete (Papadimitriou 1981), there’re lots of highly optimized heuristic solvers for MILP that provide feasible (but not guaranteed to be optimal) solutions relatively fast. In TTT we use the PuLP (https://github.com/coin-or/pulp) package and the GLPK (https://www.gnu.org/software/glpk/) solver. Note that any path through the tangle connecting the incoming and outgoing edges could yield a solution which will satisfy all constraints in (1), (2) and (4) classes. However, condition (3) may cause situations where no solution exists at all, e.g. a boundary edge could be followed by two *reasonably covered* edges that could not both be traversed at the same time. Thus, if no solution for the original problem is found, we remove the requirement that all reasonably covered edges are used to ensure that some solution will be found.

### Finding the best path with given edge multiplicities

First, we find just any path satisfying the multiplicities from the previous step. Lets replace each edge of multiplicity *n* with *n* multiedges of multiplicity 1 (and the same sequence label as the original edge) and remove all edges of multiplicity 0. Thus, the resulting problem is to find the Eulerian path in this multigraph. Since we require exactly one incoming and one outgoing edge and the tangle is connected when considered as an undirected graph, such an Eulerian path exists, and can be found using Hierholzer’s algorithm (Hierholzer and Wiener 1873).

There can be multiple traversal paths with the given multiplicities. The goal of this step is to provide the path that satisfies the multiplicities and is most consistent with the long-read alignments. We use read alignments provided by sequence-to-graph alignment tools such as GraphAligner (Rautiainen and Marschall 2020), and consider each read as a sequence of edge labels. We define allowed modifications of the path that preserve multiplicities, and use a gradient-descent-like approach, trying multiple modifications and accepting the modification only if it improves the score. For each path (which is also considered as a sequence of edge labels) its alignment score is defined as the number of reads which can be found as an exact substring in this path. Although inexact or partial pattern matching-based scoring can be more natural for this problem, the exact substring requirement allows the use of the fast Aho-Corasick algorithm (Aho and Corasick 1975) for scoring.

Let’s define a *swap* operation on a path as follows: Find two non-overlapping intervals in the path that start and end on the same vertices A and B and swap the subpaths defined by those intervals (see **Figure 4**). This operation can be considered as reordering repeat units in a repeat array.

**Figure 4.**
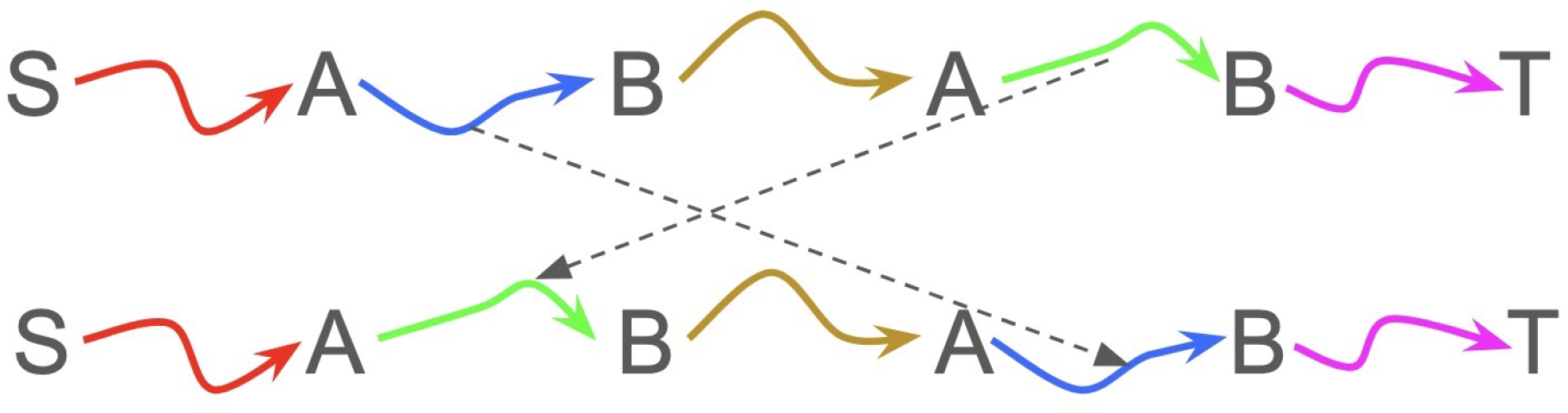
Swap operation. Capital letters represent the *vertices* of the graph. Colored arrows represent different subpaths in the assembly graph. A swap operation rearranges the green and blue subpaths.

#### Lemma 1

An Eulerian path in directed multigraph G can be transformed into any other Eulerian path in the same graph(with same start and end) through a sequence of swaps.

It is easy to see that one can iteratively extend the common prefix between two paths using swaps; by doing so, the length of the common prefix can be increased until the paths match entirely. Proof details can be found in **Supplementary Methods S1**.

TTT starts from a random Eulerian path, applies random swaps, and accepts a change only if it improves the score. The process terminates when no score-improving swap exists. This approach optimizes the consistency of read alignments to the chosen path while maintaining the edge multiplicities determined in step 1. Although Lemma 1 guarantees that there exists a sequence of swaps leading to a path with the optimal score, it is still possible for TTT to become trapped in a local optimum. To reduce this risk, we perform multiple iterations of this gradient-descent-like optimization using different random Eulerian paths as starting points. Moreover, there can be multiple solutions with exactly the same scores(i.e. in case of large exact repeat not traversed by any provided read alignment). Thus, even assuming the given coverage and assembly graph are accurate, the solution is not guaranteed to exactly match the genomic traversal but should be highly concordant with the underlying data.

### Gap resolution in zebra finch chrZ

Model sequences were aligned at bp resolution with lastal (http://last.cbrc.jp/) to the existing reference to close the gaps (https://github.com/gf777/T2T-zebra-finch/tree/main/patching). The model sequence was trimmed to include as much as possible of the sequence directly resolved by Verkko. To visualize read based evaluation, we first aligned HiFi and ONT reads to the assembly using minimap2 with the parameters `-ax map-hifi` and `-ax lr:hq` respectively. We then removed unmapped, secondary and supplementary alignments (`-F 2308`) for HiFi and standard ONT reads, and unmapped and secondary alignments (`-F 260`) for ultra-long ONT reads using SAMtools v1.22. Next, we ran NucFlag v1.0.0a2 (Vollger et al. 2019; Gao et al. 2025) by setting the window size to 50kbp for the entire chromosome, and keeping the default 5kbp for the local tangle regions. The same NucFlag workflow was applied to assess the patched regions by tool DEGAP.

### Gene models identification

To identify gene models on the resolved chromosome Z, EGAPx annotation from the T2T zebra finch assembly was transferred to the gap-resolved chromosome Z assembly using Liftoff (Shumate and Salzberg 2021). Genomic coordinates corresponding to the PAK3L cluster boundaries were defined on the resolved chromosome Z. The Liftoff-transferred annotation was intersected with these intervals to extract all gene models located within each cluster. Gene models were then screened for duplication within the cluster boundaries by identifying repeated annotations of the same gene (or gene family). Each identified protein sequence was aligned against the reference genome using miniprot (Li 2023). To simplify miniprot mapping results, we collapsed redundant alignments into a single representative call per genomic locus using an in-house script. Functional domains of the predicted proteins were identified using eggNOG-mapper v.5.0.2 (Huerta-Cepas et al. 2017). The specific analysis of PAK3L genes is documented here: https://github.com/gf777/T2T-zebra-finch/tree/main/chrZ/PAK3L.

### DEGAP analysis

DEGAP requires a left and right boundary for each gap. To match the input for TTT, we aligned the final resolved assembly to the gapped sequence and trimmed to the last aligned base before and after each gap. DEGAP v2.0 (commit 2f4c318105c77b4c72efa4d104156547977a348c) was used for all experiments. Initial runs force-joined gap2 without any inserted sequence due to a detected overlap in the boundary regions. We therefore disabled this initial merging and allowed DEGAP to use extension to fill the gap. All HiFi and ONT reads assigned to a node in the chrZ connected component by the verkko assembly were extracted. DEGAP was run with the command `DEGAP.py --mode gapfiller --seqleft $gapLeft --seqright $gapRight –hifi hifi.fastq.gz --ont ont.fastq.gz --flag left -t $cores --out $lPrefix` where `$gapLeft` was the sequence to the left of the gap and `$gapRight` was the sequencing following the gap. The `$gapRight` for gap 1 was equivalent to `$gapLeft` for gap2.

## Notes

### Competing Interest Statement

The authors have declared no competing interest.

