## Supplemental materials for "Automatic Generation of Model Sequences for Complex Regions in Assembly Graphs"

**Supplemental methods 1**:
Described methods assume that only one of *edge* and *reverse_complement(edge)* can be a part of the tangle. This is not true in general case - inverted repeats may cause both *edge* and *reverse_compement(edge)* to be a part of the same traversal.
To address this issue, following changes should be made to described methods:

a) We should distinguish the multiplicity of an edge and its reverse complement. Thus, for each edge in the assembly graph, we should have separate integer variables *mult(e)* and *mult(reverse_complement(e)).* Thus, to fix the multiple integer programming problem formulation, in condition 1 the vertex and its reverse complement should be considered separately and in conditions (2-5) *mult(e)* should be replaced by *(mult(e)* + *mult(reverse_complement(e)))*
b) Now the graph induced by the multiplicities can have more than one connected component. Because of the condition on the incoming and outgoing edges, there should be eulerian cycles through each of the components except one which contains boundary edges. Let's name those components as *cyclic components*, and the component with boundary edges as a *path component*. We can switch the orientation (replace all edges with their reverse complements) for all cyclic components without violating any of the conditions A1-A4 and changing the final score defined in A5. At least one of those new cyclic paths should be adjacent to some of the edges in the path component (otherwise the initial tangle would be disconnected), and thus the path component is extended. This can be repeated iteratively until no cyclic component remains.
c) Now we should allow not only reordering but also changing the orientation of the edges in the path. To do so, we define two additional operations: *inversion* and *reverse complement swap.*
    i) Inversion: Replace a subpath P between vertex A and its reverse complement vertex A' to its reverse complement P'
    ii) Reverse-complement swap: For two nonoverlapping intervals *(A, B)* and *(reverse_complement(B), reverse_complement(A))* between reverse-complement vertices in reverse order, reverse-complement and swap corresponding subpaths.

Those two operations can change multiplicity of an edge *e*, but preserve *(mult(e) + mult(reverse_complement(e)))*

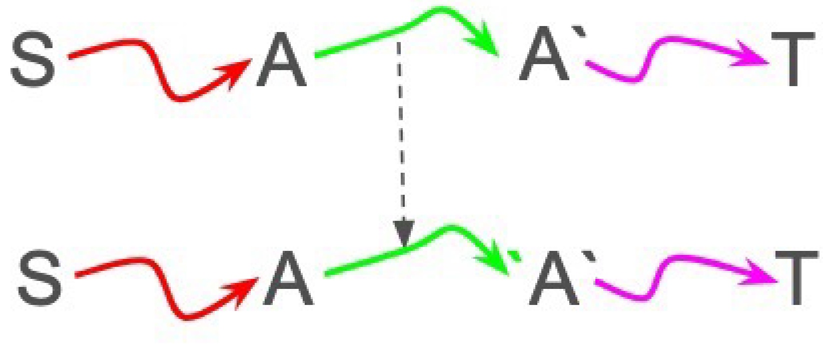

**Figure S1.** Inversion operation. Capital letters represent the vertices of the graph. Reverse-complement vertices and paths are shown with the symbol `. Inversion operation replaced the blue subpath with its reverse-complement.

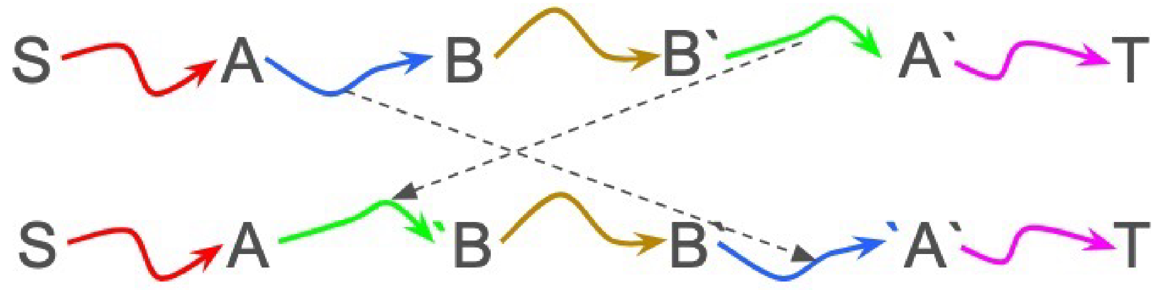


**Figure S2.** Reverse-complement swap operation. Capital letters represent the vertices of the graph. Reverse-complement vertices and paths are shown with the symbol `. Blue and green subpaths are replaced by their reverse-complements and swapped.

To process tangles with two traversing haplotype paths (from S1 to T1 and from S2 to T2) after determining edge multiplicities we add auxiliary edge *aux* of multiplicity 1 from the end of first traversal T1 to start of the second path S2. Then, the final path is split in two by removing *aux* from it. To preserve pairing of incoming/outgoing edges, during optimization step C we forbid inversion of intervals containing *aux.*

**Lemma 1:** An Eulerian path in directed graph G can be transformed into any other Eulerian path in the same graph(with same start and end) through a sequence of swaps.

Let's proceed by contradiction. Assume there exist at least two distinct Eulerian paths from a starting vertex *S* to a terminal vertex *T* in a graph *G*. Let $P_1$ and $P_2$ be two such paths, with the premise that $P_1$ cannot be transformed into $P_2$ through any sequence of path swaps.

Let Ω be the set of all paths that can be generated by applying a series of swaps to $P_1$. From this set Ω, let's select a path, which we'll call *P′*, that has the longest possible common prefix with $P_2$. Our goal is to show that if *P′* is not identical to $P_2$, we can always perform a swap on *P′* to create a new path, *P″*, which has an even longer common prefix with $P_2$. This will contradict the initial choice of *P′* as the path with the maximal common prefix.

Let the maximal common prefix of P' and $P_2$ be denoted by $P_{prefix}$, which terminates at a vertex *X*. Let the first edge after vertex *X* in path *P′* be *a*, and in path $P_2$ be *b*. By the definition of a maximal prefix, it must be that $a \neq b$.

Let us now consider a subgraph, $G_r$, formed by removing all edges of $P_{prefix}$ from the original graph $G$. The remaining portions of the paths, $P'_r$ (from $P'$) and $P_{2r}$ (from $P_2$), are both Eulerian paths within $G_r$ , each starting at vertex $X$ and ending at vertex $T$. The path $P'_r$ begins with edge $a$, while $P_{2r}$ begins with edge $b$. Since $P'_r$ is an Eulerian path in $G_r$, it must traverse every edge in $G_r$ , including edge $b$. Because $P_{2r}$ starts at $X$ with edge $b$, the source of edge $b$ must be $X$.

Now our goal is to find a swap that would transform $P'_r$ to start with b. If we find it, the same swap would increase the common prefix length between P' and $P_2$ which contradicts the choice of P'. Therefore, we can decompose the path $P'_r$ into three distinct segments:

1. A cycle, $\mathbf{P'_c}$ , which starts at $X$ with edge $a$ and eventually returns to $X$.
2. The single edge, $b$ which connects $X$ to a vertex we'll call $Z$.
3. A suffix path, $\mathbf{P'_s}$ , which runs from vertex $Z$ to the terminal vertex $T$.

Now, we will show that the vertex sets of $P'_c$ and $P'_s$ must be disjoint. Suppose they are not. This means there exists some vertex $Y$ that is common to both the cycle $P'_c$ and the path $P'_s$. Then we could then perform a swap of X-Y fragments in $P'_c$ and $(b \rightarrow P'_s)$, and the resulting path would start from b. Let $V'_c$ be the set of vertices in $P'_c$, and let $V'_s$ be the set of vertices in $P'_s$. We have established that $V'_c \cap V'_s = \varnothing$.

Finally, consider the path $P_{2r}$. It starts from $X$ with edge $b$ and, as an Eulerian path, must traverse all edges of $Gr$, which includes the edge a starting from vertex X. This implies there must be a path from $X \in V'_s$ to $X \in V'_c$. Consequently, there must exist an edge, let's call it $e = (A, B)$, such that its source vertex $A \in V'_s$ and its destination vertex $B \in V'_c$ .

Where could this edge $e$ possibly lie in the path $P'_r = P'_c \rightarrow b \rightarrow P'_s$?

- It cannot be in $P'_c$, because its source vertex $A$ is not in $V'_c$.
- It cannot be in $P'_s$, because its destination vertex $B$ is not in $V'_s$.
- It's not $b$ since destination of $b$ is in $V'_s$

The edge $e$ exists in the graph $G_r$ but is not traversed by the path $P'_r$. This is a terminal contradiction, as it violates the premise that $P'$ is an Eulerian path. Thus, our initial assumption must be false and any Eulerian path can be transformed into any other through a series of swaps.

**Supplemental methods 2**:
Precisely extracting tangles from an assembly graph is a complex question out of scope of this work. To minimize arbitrary decisions, we designed the following process for tangle identification based on the input assembly graph

1. Remove all bubbles (defined as an edge pair that shares incoming and outgoing vertices and have similar lengths - $l_1 < l_2 * 1.5;\ l_2 < l_1 * 1.5$)
2. Remove all tips (defined as edges with either no outgoing or incoming connections and shorter than 30kbp)
3. Compress all nonbranching paths

4. Repeat steps 1-3 until no further simplification is possible. Remove all edges that are longer than 1 Mbp
6. Consider all connected components in the remaining graph as tangle candidates
7. For each tangle candidate iteratively restore all adjacent edges that were removed in steps 1-4
8. Consider all edges adjacent to the tangle (in the initial graph) as boundary edges. If a boundary edge has more than one adjacent incoming(outgoing) edge and those edges are not in the tangle - add that boundary edge to the tangle and consider those incomings/outgoings as boundaries.

The aim of this pipeline is to get rid of large but trivial to resolve haploid and diploid regions and to reduce the regions in consideration as much as possible. Although TTT can be run on large tangles, an underlying assumption of coverage uniformity across the tangle is often violated for multimegabase-scale regions.

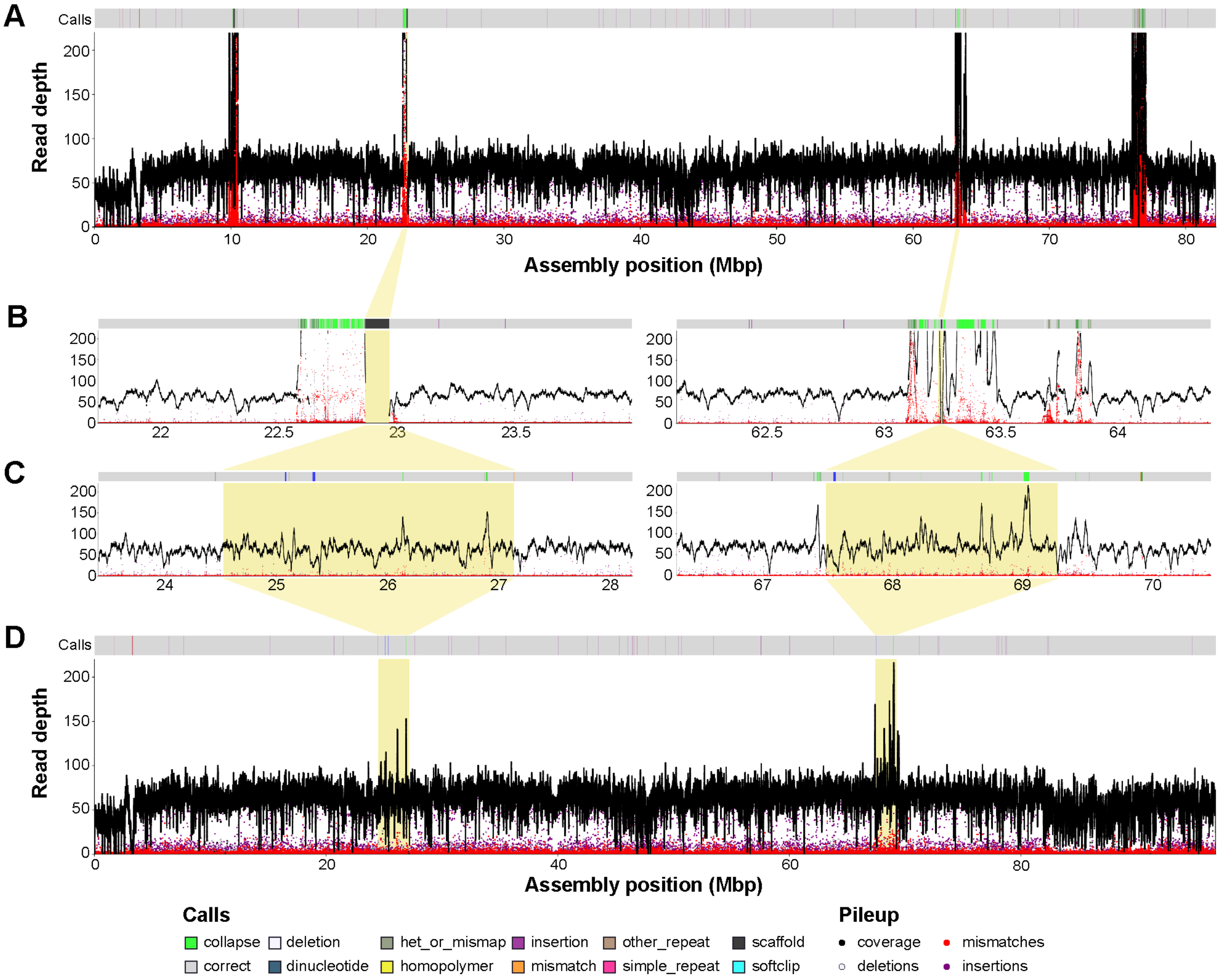

**Figure S3.** Nucflag HiFi coverage plots for (A) zebra finch chromosome Z before resolving the gaps (NCBI accession NC_133063.1), two tangle regions (B) before and (C) after running TTT, and (D) zebra finch chromosome Z after resolving the four gaps. The start and end position for the plotting region in (B)

and (C) are the same for each tangle, with the tangles highlighted in yellow. The total collapsed regions decreased from 124,253bp to 19,908bp in tangle 1 and from 147,035bp to 71,869bp in tangle 2.

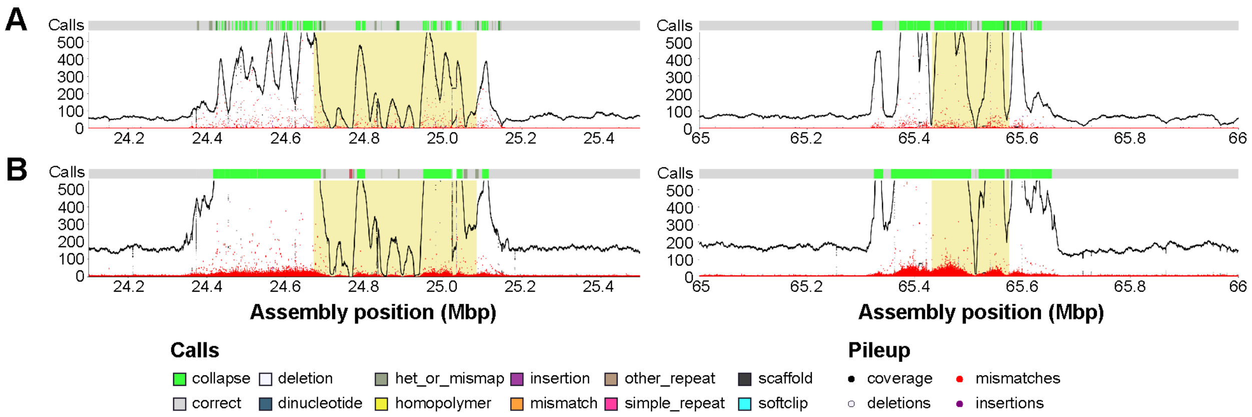


**Figure S4.** Nucflag plots based on (A) HiFi reads and (B) ONT reads for two tangle regions patched using DEGAP. Left is tangle 1 and right is tangle 2. The two tangle regions remain unresolved, with collapsed reads shown in green.

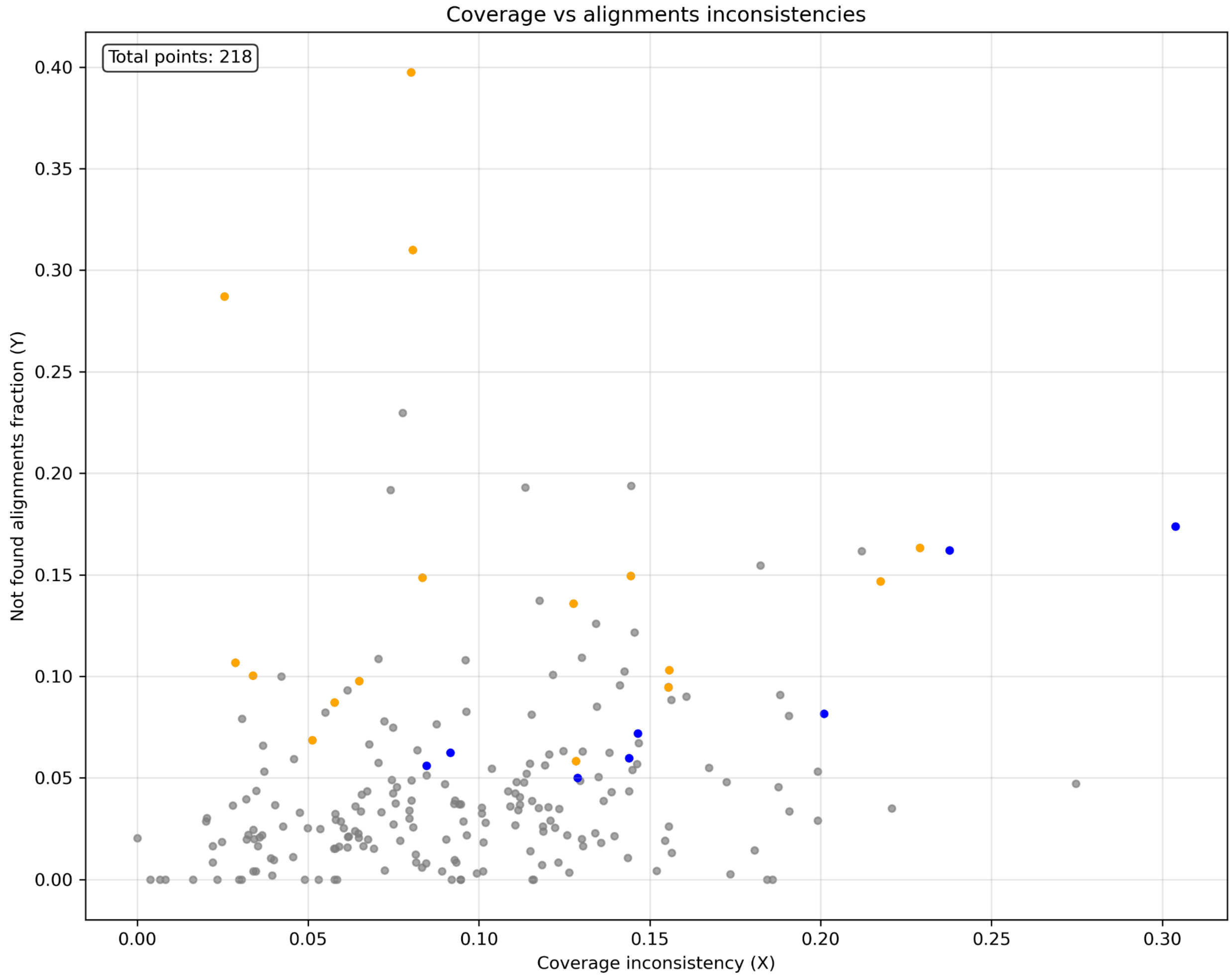


**Figure S5.** Internal TTT evaluation on tangles of the HG002 hifi-only assembly. Orange points represent the tangles where verkko's resolution is better than TTT's and blue points represent the tangles where TTT provides better resolution.
